# Slice’N’Dice: Maximising the value of predicted models for structural biologists

**DOI:** 10.1101/2022.06.30.497974

**Authors:** Adam J. Simpkin, Luc G. Elliott, Kyle Stevenson, Eugene Krissinel, Daniel J. Rigden, Ronan M. Keegan

## Abstract

With the advent of next generation modelling methods, such as *AlphaFold2*, structural biologists are increasingly using predicted structures as search models for Molecular Replacement (MR) when experimental structures of homologues are unavailable. Inaccuracy in domain-domain orientations is often a key limitation when using predicted models for MR. *Slice’N’Dice* is a software package designed to address this issue by first slicing models into distinct structural units and then automatically placing the slices using *Phaser*. The slicing step can use *AlphaFold2*’s predicted aligned error (PAE), or can operate via a variety of Cα atom clustering algorithms, extending applicability to structures of any origin. The number of splits can be selected by the user. *Slice’N’Dice* is available in CCP4 8.0 and is currently being adapted for cryo-EM use cases.

## 1 Introduction

Molecular replacement (MR) remains the dominant method for solving the phase problem in Macromolecular X-ray crystallography (MX) with 87.2% of the crystal structures deposited in the Protein Data Bank (PDB) (Burley *et al*., 2021) having been solved by MR from June 2021 to June 2022. The emergence of next generation predicted models has wide-reaching implications for MX with MR being a key application. The availability of sufficiently close homologues with experimentally determined structures has always been a limitation in MR, one which is largely solved by the highly accurate models produced by next generation modelling methods, such as *AlphaFold2* (Jumper *et al*., 2021) and *RosettaFold* (Baek *et al*., 2021).

Despite the high quality of predicted models, some preprocessing is often required for success in MR. The quality of the predicted model varies across the target sequence with some regions being inaccurately predicted. Both *AlphaFold2* and *RosettaFold* provide predicted quality scores on a per-residue basis that can be used to assess the accuracy of the predicted models and guide the removal of any residues likely to have been inaccurately modelled. *AlphaFold2* gives the predicted Local Distance Difference Test (pLDDT) score (Jumper *et al*., 2021), a per-residue estimate of its confidence on a scale from 0-100 where higher values correspond to higher confidence and *RosettaFold* gives an estimated Root Mean Square Deviation (RMSD) score, a per-residue estimate of the RMSD to the true structure where lower values correspond to higher confidence. Both methods store this information in the B-factor column of their output PDB files. For PDB-derived search models, B-factors are used for weighting search models in *Phaser (McCoy et al., 2007)*, and therefore converting pLDDT and RMSD values to predicted B-factors can improve the performance of the models in MR (Croll *et al*., 2019). While confidence scores work well for assessing the reliability of individual residues, they are unable to indicate global inaccuracies in the model such as those caused by inter-domain conformational changes. *AlphaFold2* provides a Predicted Aligned Error (PAE) (Varadi *et al*., 2022) matrix which can be used to assess the reliability of a given prediction of inter-domain orientation.

Here, we present *Slice’N’Dice*, an automated MR pipeline designed to pre-process predicted models by removing low confidence regions and converting confidence scores into predicted B-factors. *Slice’N’Dice* will also slice predicted models into distinct structural units which are then placed in an automatic MR pipeline. These steps allow *Slice’N’Dice* to maximise the effectiveness of predicted models in MR. Current work is focussing on adapting *Slice’N’Dice* to CryoEM use cases.

## 2 Methods

*Slice’N’Dice* is a combination of two methods: ‘Slice’ which breaks models up into distinct structural units and ‘Dice’ which performs MR if a file containing crystal diffraction data is provided.

### 2.1 Slice

#### 2.1.1 Clustering

Clustering algorithms are used to detect distinct structural units within a predicted model. *Slice’N’Dice* provides eight clustering methods for users to choose from (Fig. 1). The methods from *SciKitLearn* (Garreta & Moncecchi, 2013) cluster based on the coordinates of the Cα atoms, whereas the methods from *CCTBX* cluster on the Predicted Aligned Error (PAE) produced by *AlphaFold2* (Baek et al., 2021). Based on preliminary data, the Birch algorithm is the current default but further benchmarking will be done.

**Figure 1).**
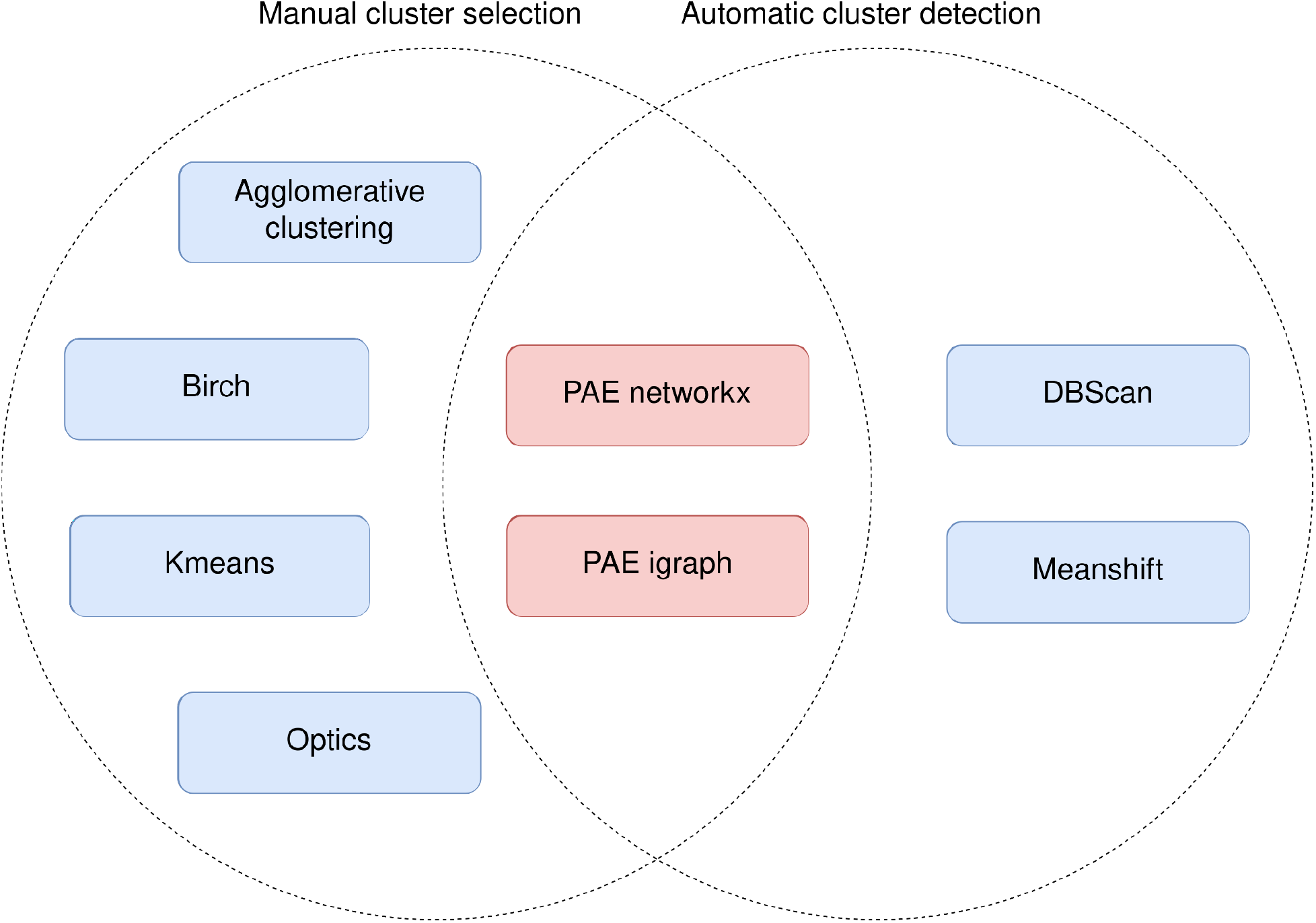
Venn diagram showing the various clustering methods included in *Slice’N’Dice*. Shown in blue are clustering methods included in *SciKitLearn* that cluster based on Cα atom coordinates. Shown in red are clustering methods included in *CCTBX* that cluster based on the Predicted Aligned Error (PAE) from *AlphaFold2*. On the left are all the clustering methods that require the number of clusters to be specified and on the right are clustering methods that automatically determine the number of clusters. *CCTBX*’s PAE methods automatically identify clusters, however if a user defines a maximum number of splits, the closest clusters are merged until the number of splits ≤ maximum number of splits.

**Figure 2).**
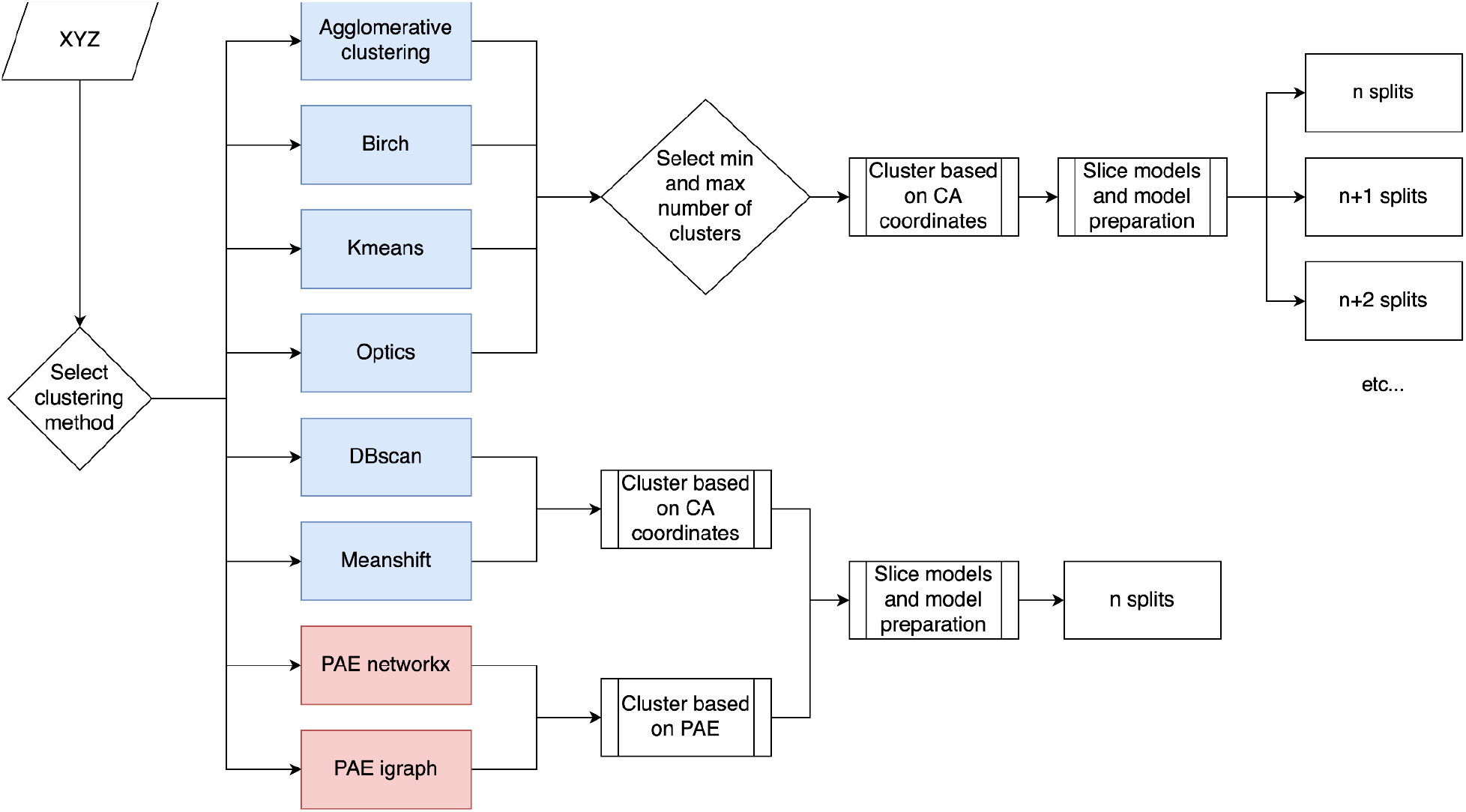
Flowchart showing the model slicing process. SciKitLearn methods are shown in blue and CCTBX methods are shown in red.

The clustering methods can be subdivided further into those methods which automatically determine the number of clusters to produce and those methods which require users to manually specify the number of clusters to produce. For those methods where the number of clusters needs to be specified, users can provide *Slice’N’Dice* with the `min_splits` and `max_splits` arguments. This allows slice and dice to test a range of different splits.

If, for example, a target protein is very large and needs to be modelled in several chunks, the `xyz_list` argument can be used to provide *Slice’N’Dice* with several input models. Additionally the `xyz_list_splits` argument can be used to specify the number of times to split each input model if using a clustering method that requires manual selection of the number of clusters.

#### 2.1.2 Model preparation

The `xyz_source` option can be used to specify the origin of a model (e.g. *AlphaFold2, RosettaFold*). Predicted models often require preparation to succeed in MR. In *Slice’N’Dice* we do this in two ways:

1. Per residue quality scores (e.g. pLDDT) recorded in the B-factor column of the PDB file are converted to predicted B-factors using the methods described in Simpkin *et al. (Simpkin et al., *2022*)* and Croll *et al. (Croll et al., 2019)*.
2. Residues that fall below a threshold score are removed from the model.

For *AlphaFold2* models, the default pLDDT threshold is 70 and can be set at the command line with the `plddt_threshold` flag and for *RosettaFold* models, the default RMS threshold is 1.75 and can be set at the command line with the `rms_threshold` flag. For *AlphaFold2* models where the pLDDT scores have already been converted into predicted B-factors, the `xyz_source` can be set to `alphafold_bfactor`. If `alphafold_bfactor is` used in combination with the `plddt_threshold` flag, the threshold is first converted into a predicted B-factor and any residues above this value are removed.

### 2.2 Dice

The second part of the *Slice’N’Dice* pipeline, ‘Dice’, performs molecular replacement on the individual slices produced by ‘Slice’. In simple mode, ‘Dice’ provides all the slices to *Phaser* (McCoy *et al*., 2007) simultaneously to automatically place as many slices as possible. In intensive mode, ‘Dice’ first attempts to place the slices with *Phaser* individually. Any slices which appear to have been placed correctly (LLG ≥ 60) are then combined into a single *Phaser* job using one of the solutions as a fixed model to ensure that they are all at the same origin. The result of this combined *Phaser* job is used as an initial fixed model for further MR steps.

*MOLREP’s* Phased Translation Function (PTF) mode (Vagin & Teplyakov, 2010), which makes use of the phase information given by the initial fixed model to assist in MR, is then used to try and place the slices that could not be positioned in the initial *Phaser* job. After each *MOLREP* job, *REFMAC5* (Murshudov *et al*., 2011) is used to assess if the placed slice has improved the solution. Figure 3 shows the decision-making process used in the intensive mode.

**Figure 3).**
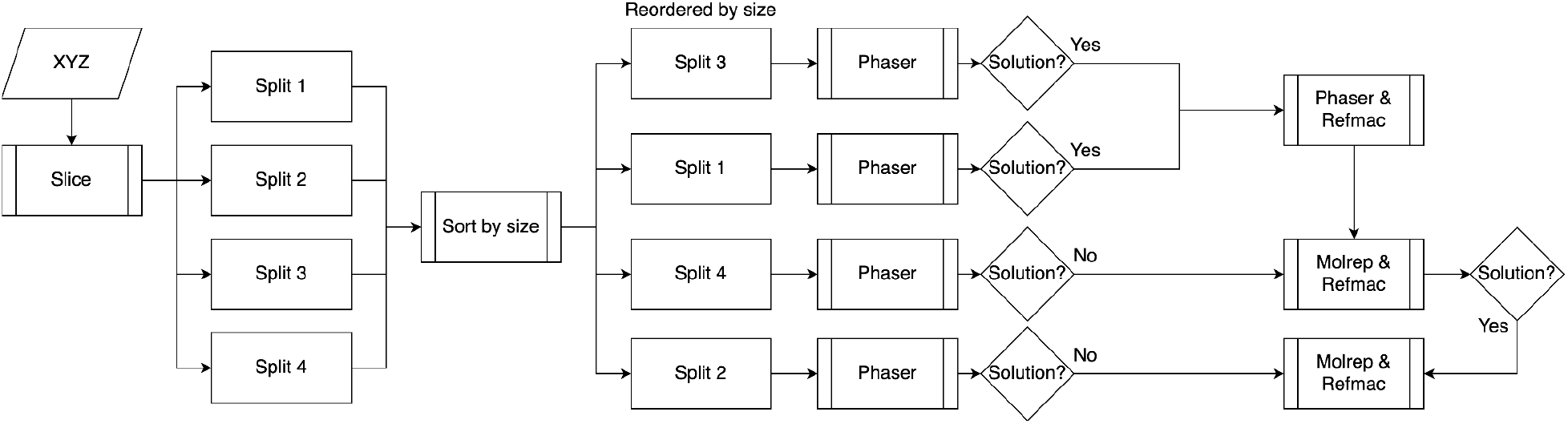
Flowchart showing *Slice’N’Dice*’s intensive MR mode.

## 3 Results

*Slice’N’Dice* can significantly improve the performance of structure predictions used as search models for MR, solving cases that would otherwise be difficult or intractable. Here we show a number of examples that highlight the ways in which *Slice’N’Dice* can maximise the effectiveness of predicted models in MR.

### 3.1 Example 1 - 7OA7

7OA7 is a crystal structure of a PilC minor pilin solved by Single-wavelength Anomalous Dispersion (SAD). At the time of its release, the closest hit in the PDB, 3ASI, only has 12% sequence identity to the target and was insufficiently similar to work in MR (Fig. 4A). A model made by *AlphaFold2* was very good quality overall (average pLDDT: 85.61), however was unable to solve the structure as *AlphaFold2* modelled a different conformation between the two domains (Fig. 4B). By using the *Birch* algorithm in *Slice’N’Dice* to split the structure into two, the structure can readily be solved by MR (Table 1, Fig. 4C). This structure could also be solved using the PAE *networkx* algorithm (Hagberg *et al*., 2008) with the maximum number of splits set to two (Table 1). *Birch* and PAE *networkx* identified slightly different domain boundaries (Suppl. Fig. 1), and whilst *Birch* seemed to work slightly better in this case, both methods could be refined to the same point.

**Table 1).** Summary of the example results. For 7OA7 the results using PAE *networkx* are shown in brackets.

|  | 7OA7 | 7RB4 | 7B9C |
| --- | --- | --- | --- |
| Resolution (Å) | 1.45 | 2.19 | 2.4 |
| No. copies ASU | 1 | 1 | 1 |
| Clustering method | *Birch* (PAE *networkx*) | *Birch* | *Birch* |
| Number of slices | 2 | 3 | 4 |
| RMSD to target before *Slice'N'Dice* | 1.528 (1.083) | 3.187 | 4.236 |

| (Å) |  |  |  |
| --- | --- | --- | --- |
| RMSD to target after *Slice'N'Dice* (Å) | 0.505, 0.550 (1.213, 0.488) | 0.528, 0.757, 0.9 | 0.466, 0.616, 0.797, 1.038 |
| LLG | 1339 (641) | 114 | 310 |
| TFZ | 35.6 (18.5) | 13.2 | 24.1 |
| R-factor | 0.41 (0.41) | 0.44 | 0.48 |
| R-free | 0.41 (0.41) | 0.47 | 0.51 |
| Local Map CC | 0.79 (0.79) | 0.72 | 0.80 |
| Global Map CC | 0.7 (0.7) | 0.59 | 0.47 |

**Figure 4).**
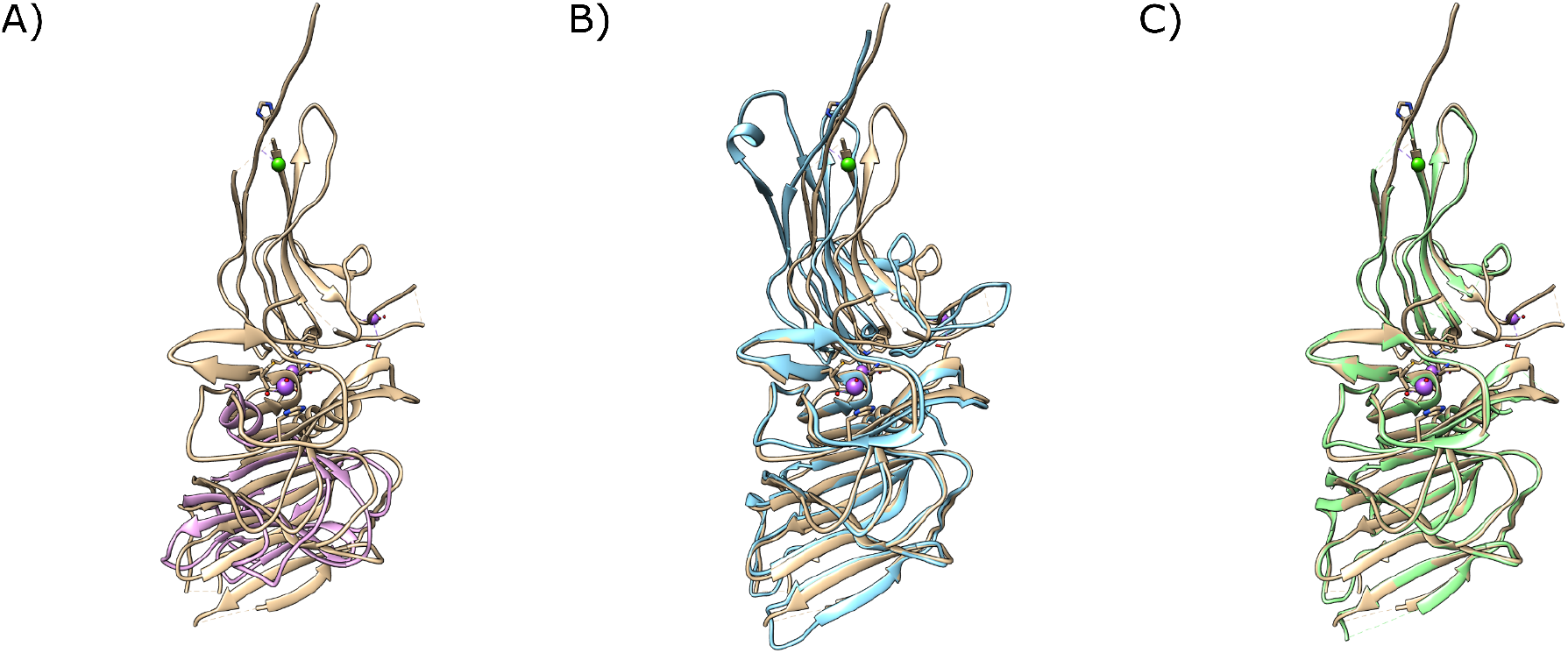
A - The closest match in the PDB to the target structure, 3ASI (pink), superimposed on the crystal structure of 7OA7 (tan). B - An *AlphaFold2* model of the target (blue) superimposed on the crystal structure of 7OA7 (tan). C - The *AlphaFold2* model after slicing and MR with *Slice’N’Dice* (green) shown against the crystal structure of 7OA7 (tan). Figure made using UCSF Chimera (Pettersen *et al*., 2004).

### 3.2 Example 2 - 7RB4

7RB4 is a crystal structure of Peptono Toxin solved by SAD. The closest hit in the PDB, 1F0L only has 26% sequence identity to the target and was insufficiently similar to work in MR, even when split with *Slice’N’Dice* (Fig. 5A). A model made by *AlphaFold2* was poor quality overall (average pLDDT: 61.02, Fig. 5B). Indeed, only splitting the model with *Slice’N’Dice* failed to lead to a structure solution. Nonetheless, the combination of removal of residues below a relaxed pLDDT threshold of 50 with splitting the model into three units, steps implemented together in *Slice’N’Dice*, led to structure solution (R-fact: 0.44, R-free: 0.47, mapCC: 0.59, Fig. 5C). This solution could be significantly improved by running 20 cycles of *Buccaneer* (Cowtan, 2006) which increased the percentage of modelled residues from 34 to 74 (Completeness by residues: 0.74, R-fact: 0.23, R-free: 0.30, Global mapCC: 0.84, Fig. 5D).

**Figure 5).**
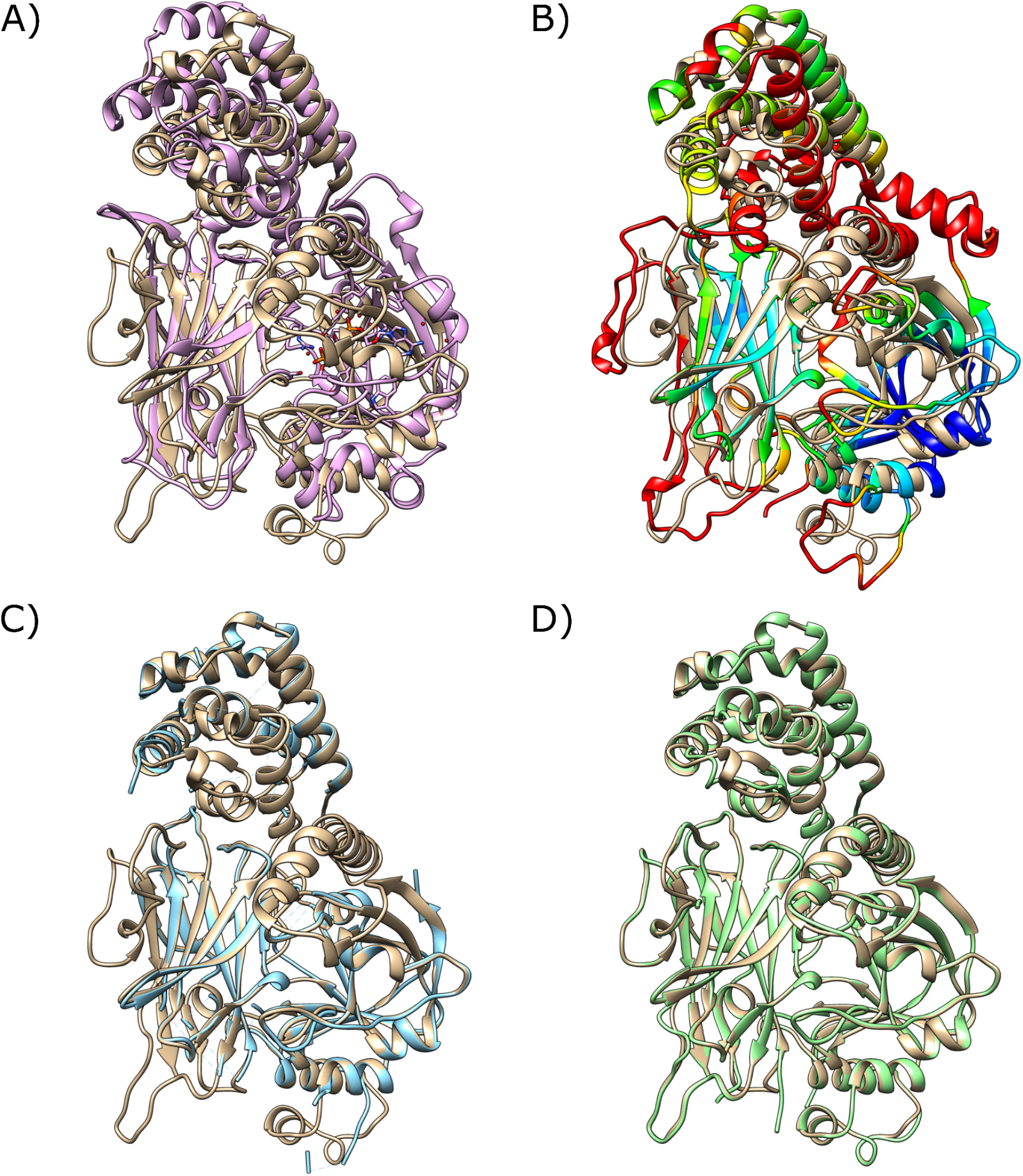
A - The closest match in the PDB to the target structure, 1F0L (pink), superimposed on the crystal structure of 7RB4 (tan). B - An *AlphaFold2* model of the target coloured on a scale of red_yellow_green_cyan_blue where red indicates a low pLDDT score (≤50) and blue indicates a high pLDDT score (≥90), superimposed on the crystal structure of 7RB4 (tan). C - The *AlphaFold2* model after preprocessing, slicing and MR with *Slice’N’Dice* (blue), shown against the crystal structure of 7RB4 (tan). D - The placed *AlphaFold2* model after 20 cycles of model building with *Buccaneer* (green) shown against the crystal structure of 7RB4 (tan). Figure made using UCSF Chimera (Pettersen *et al*., 2004).

### 3.3 Example 3 - 7B9C

7B9C is a crystal structure of a minimal Splicing factor 3B (SF3B) core in complex with spliceostatin A solved by MR using PDBs 5IFE and 6EN4 as search models. Despite highly similar homologues in the PDB, a model of a SF3B subunit one deposited in the EBI *AlphaFold* Protein Structure Database (Varadi *et al*., 2022) (Uniprot ID: O75533) was insufficiently similar to the target protein to succeed in MR (Fig 6A). The HEAT repeat region of SF3B is confidently predicted by *AlphaFold2* but has been predicted to adopt a much tighter conformation than the crystal structure. Without reference to the solved structure, it would be unclear to the experimentalist where the model should be split manually in order for it to succeed in MR. However, *Slice’N’Dice*’s automated slicing procedure was able to successfully slice the model into four structural units of which three could be placed by MR and used to solve the structure (Fig. 6B). The refinement scores were a little high (R-fact: 0.48, R-free: 0.51) due to the fact that the SF3B subunit one domain made up only 43.8% of the total scattering content. Nonetheless, this could be confirmed as a true solution using *phenix*.*get_cc_mtz_pdb* (Liebschner *et al*., 2019) (Global mapCC: 0.47).

**Figure 6).**
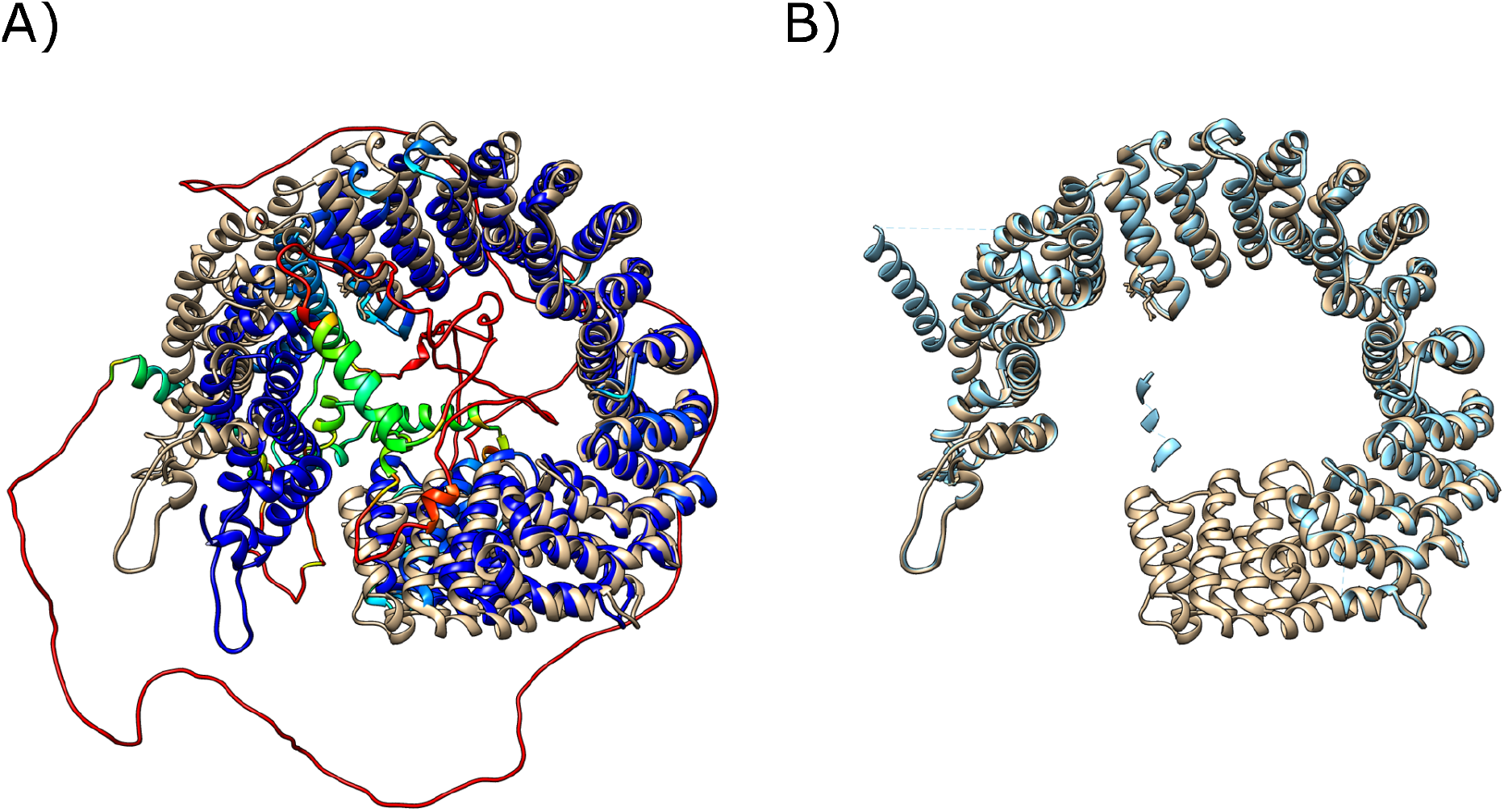
A - O75533, a model from the EBI *AlphaFold2* database coloured on a scale of red_yellow_green_cyan_blue where red indicates a low pLDDT score (≤50) and blue indicates a high pLDDT score (≥90), aligned to the SF3B core (chain C) from 7B9C (tan). B - O75533 after preprocessing, slicing and MR with *Slice’N’Dice* (blue), shown against the SF3B core (chain C) from 7B9C (tan). Figure made using UCSF Chimera (Pettersen *et al*., 2004).

**Figure 6).**
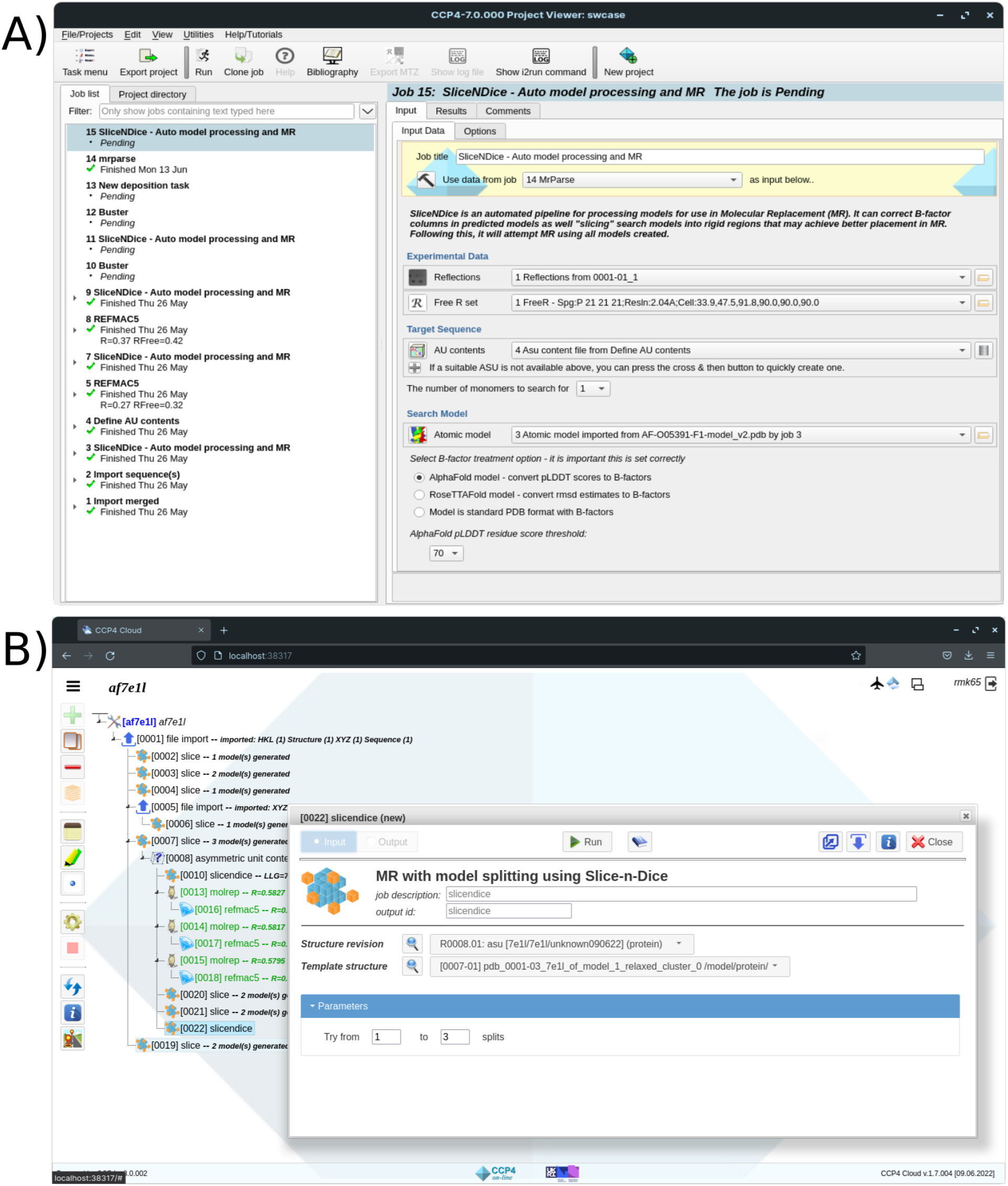
A - CCP4i2 interface page for *Slice’N’Dice*. B - CCP4-Cloud interface page for *Slice’N’Dice*.

## 4 GUIs

*Slice’N’Dice* has been incorporated into several CCP4 graphical user interfaces (GUIs) including CCP4i2 and CCP4-Cloud. These provide interfaces for slicing up models without performing MR (*Slice*) and slicing up the models with automated MR (*Slice’N’Dice*).

Running *Slice’N’Dice* on the command line will still allow users to select more run options, however the CCP4 interfaces provide a quick and easy way to run *Slice’N’Dice. Slice’N’Dice* GUIs are currently being developed for CCP4-online and CCPEM.

## 5 Discussion and Conclusions

*Slice’N’Dice* offers crystallographers an easy and automated means to address cases where the conformation of a structure prediction, especially in terms of inter-domain orientations, differs significantly from that of the target. In such cases *Slice’N’Dice* can significantly improve the chance of MR success. Here we showed that clustering algorithms can be used to identify distinct structural units within a model that may not be immediately obvious when visually inspecting the structure. Currently, the default clustering algorithm used by *Slice’N’Dice* is Birch (Zhang *et al*., 1996). While Birch has performed well through the development stage of *Slice’N’Dice*, large scale testing will now be undertaken to identify which clustering algorithm performs best. Future work might additionally look into the inclusion of other clustering methods such as *SWORD2* (Cretin *et al*., 2022) and *DCI* (Kumar *et al*., 2022), and the combination of clustering methods in a consensus strategy. We’re also aware that clustering in combination with the removal of low confidence residues can occasionally leave disconnected fragments (Fig. 6B): recognising that these might impact on packing of solutions, we will explore methods to identify and eliminate these. The latter, using predicted motions for definition of structural units, might be particularly relevant given that the dynamic properties of multi-domain proteins underlie some of the difficulties that *Slice’N’Dice* is designed to address. Finally, recognising that predicted models are increasingly being used to solve macromolecular structures in CryoEM (Kryshtafovych *et al*., 2021; Slavin *et al*., 2021; Ivanov *et al*., 2022; Skalidis *et al*., 2022), *Slice’N’Dice* is currently being developed for CryoEM use cases where the slicing algorithm will be partnered with an automated map fitting pipeline.

