## Supplemental figure 1 for "Slice’N’Dice: Maximising the value of predicted models for structural biologists"

### Supplementary

A) B)

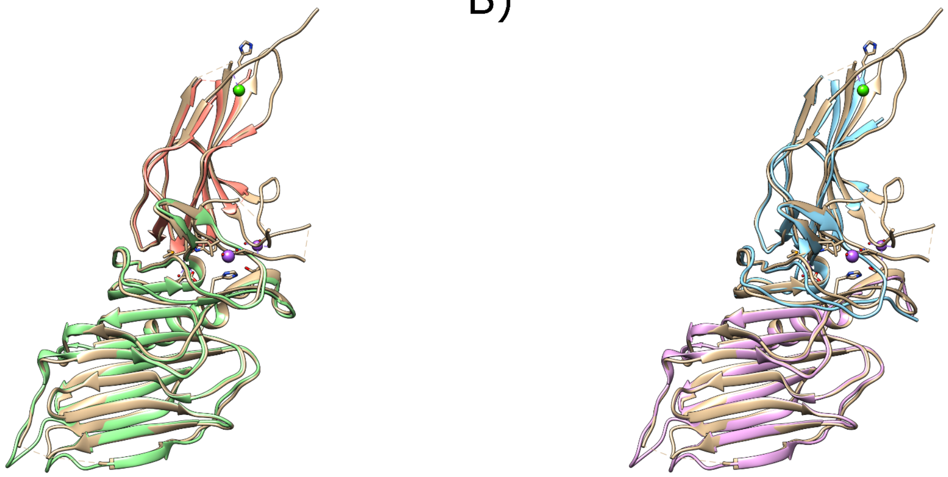

Supplementary Figure 1) A) Slices made using the *Birch* algorithm (orange, green) superimposed on the crystal structure of 7OA7 (tan). B) Slices made using the PAE *networkx* algorithm (blue, pink) superimposed on the crystal structure of 7OA7 (tan).
